# Carbon monoxide dehydrogenases enhance bacterial survival by oxidising atmospheric CO

**DOI:** 10.1101/628081

**Authors:** Paul R.F. Cordero, Katherine Bayly, Pok Man Leung, Cheng Huang, Zahra F. Islam, Ralf B. Schittenhelm, Gary M. King, Chris Greening

**Affiliations:** School of Biological Sciences, Monash University, Clayton, VIC 3800, Australia; Monash Biomedical Proteomics Facility and Department of Biochemistry, Monash Biomedicine Discovery Institute, Monash University, Clayton, VIC 3800, Australia; School of Biological Sciences, Louisiana State University, Baton Rouge, LA 70803, LA USA

## Abstract

Carbon monoxide (CO) is a ubiquitous atmospheric trace gas produced by natural and anthropogenic sources. Some aerobic bacteria can oxidize atmospheric CO and, collectively, they account for the net loss of ~250 teragrams of CO from the atmosphere each year. However, the physiological role, genetic basis, and ecological distribution of this process remain incompletely resolved. In this work, we addressed these knowledge gaps through culture-based and culture-independent work. We confirmed through shotgun proteomic and transcriptional analysis that the genetically tractable aerobic soil actinobacterium *Mycobacterium smegmatis* upregulates expression of a carbon monoxide dehydrogenase by 50-fold when exhausted for organic carbon substrates. Whole-cell biochemical assays in wild-type and mutant backgrounds confirmed that this organism aerobically respires CO, including at sub-atmospheric concentrations, using the enzyme. Contrary to current paradigms on CO oxidation, the enzyme did not support chemolithoautotrophic growth and was dispensable for CO detoxification. However, it significantly enhanced long-term survival, suggesting that atmospheric CO serves a supplemental energy source during organic carbon starvation. Phylogenetic analysis indicated that atmospheric CO oxidation is widespread and an ancestral trait of CO dehydrogenases. Homologous enzymes are encoded by 685 sequenced species of bacteria and archaea, including from seven dominant soil phyla, and we confirmed genes encoding this enzyme are abundant and expressed in terrestrial and marine environments. On this basis, we propose a new survival-centric model for the evolution of CO oxidation and conclude that, like atmospheric H_2_, atmospheric CO is a major energy source supporting persistence of aerobic heterotrophic bacteria in deprived or changeable environments.

## Introduction

Carbon monoxide (CO) is a chemically reactive trace gas that is produced through natural processes and anthropogenic pollution. The average global mixing ratio of this gas is approximately 90 ppbv in the troposphere (lower atmosphere), though this concentration greatly varies across time and space, with levels particularly high in urban areas [1–4]. Currently, human activity is responsible for approximately 60% of emissions, with the remainder attributable to natural processes [1]. Counteracting these emissions, CO is rapidly removed from the atmosphere (lifetime of two months) by two major processes: geochemical oxidation by atmospheric hydroxyl radicals (85%) and biological oxidation by soil microorganisms (10%) [1, 5]. Soil microorganisms account for the net consumption of approximately 250 teragrams of atmospheric CO [1, 5, 6]; on a molar basis, this amount is seven times higher than the amount of methane consumed by soil bacteria [7]. Aerobic CO-oxidizing microorganisms are also abundant in the oceans; while oceans are a minor source of atmospheric CO overall [8, 9], this reflects that substantial amounts of the gas are produced photochemically within the water column and the majority is oxidized by marine bacteria before it is emitted to the atmosphere [10].

Aerobic CO-oxidizing microorganisms can be categorized into two major groups, the carboxydotrophs and carboxydovores [11]. The better studied of the two groups, carboxydotrophs grow chemolithoautotrophically with CO as the sole energy and carbon source when present at elevated concentrations. To date, this process has been reported in 11 bacterial genera from four classes **(Table S1)**: Alphaproteobacteria [12–15], Gammaproteobacteria [12, 15–18], Actinobacteria [19–21], and Bacilli [22]. Genetic and biochemical studies on the model alphaproteobacterial carboxydotroph *Oligotropha carboxidivorans* have demonstrated that form I carbon monoxide dehydrogenases mediate aerobic CO oxidation [23–25]. The catalytic subunit of this heterotrimeric enzyme (CoxL) contains a molybdenum-copper center that specifically binds and hydroxylates CO [24, 25]. In such organisms, electrons derived from CO oxidation are relayed through both the aerobic respiratory chain to support ATP generation and the Calvin-Benson cycle to support CO_2_ fixation [11, 26]. With some exceptions [19], these CO dehydrogenases have a high catalytic rate but exhibit low-affinity for their substrate (*K*_m_ > 400 nM) [27]. Thus, carboxydotrophs can grow in specific environments with elevated CO concentrations, but often cannot oxidize atmospheric CO [11, 28].

Carboxydovores are a broader group of bacteria and archaea adapted to oxidize CO at lower concentrations, including atmospheric levels, in a broad range of environments. These bacteria can oxidize CO but, in contrast to carboxydotrophs, require organic carbon for growth [11, 29]. Carboxydovores have now been cultured from some 31 bacterial and archaeal genera to date **(Table S1)**, spanning classes Alphaproteobacteria [29–32], Gammaproteobacteria [29, 33–36], Actinobacteria [18, 37–40], Bacilli [41], Thermomicrobia [41–44], Ktedonobacteria [44, 45], Deinococcota [41], Thermoprotei [46, 47], and Halobacteria [33, 48]. Carboxydovores are also thought to use form I CO dehydrogenases, but usually encode slower-acting, higher-affinity enzymes. In contrast to carboxydotrophs, carboxydovores usually lack a complete Calvin-Benson cycle, suggesting they can support aerobic respiration, but not carbon fixation, using CO [11]. A related enzyme family (tentatively annotated as form II CO dehydrogenases) was also proposed to mediate CO oxidation in carboxydovores [11, 29, 49], but recent studies suggest CO is not their physiological substrate [32].

The physiological role of CO oxidation in carboxydovores has remained unclear. It was originally thought that such microorganisms oxidize CO primarily to support mixotrophic growth [29, 30], but a recent study focused on the alphaproteobacterial carboxydovore *Ruegeria pomeroyi* showed that CO neither stimulated growth nor influenced metabolite profiles [31]. We recently developed an alternative explanation: consumption of atmospheric CO enables carboxydovores to survive carbon limitation [44, 50, 51]. This hypothesis is inspired by studies showing atmospheric H_2_ oxidation enhances survival [44, 52–57]. In support of this, CO dehydrogenases have been shown to be upregulated by five different bacteria during carbon limitation [38, 44, 53, 58, 59] and atmospheric CO is consumed by stationary-phase cells [44, 60]. Moreover, ecological studies have shown that CO is rapidly oxidized in ecosystems containing low organic carbon [51, 61, 62]. However, in contrast to atmospheric H_2_ [53–55, 57, 63], it has not yet been genetically or biochemically proven that atmospheric CO supports survival. To address this, we studied CO oxidation in *Mycobacterium smegmatis*, a genetically tractable representative of a globally abundant soil actinobacterial genus [64, 65]. We show, through proteomic, genetic, and biochemical analyses, that a form I CO dehydrogenase is (i) strongly induced by organic carbon starvation, (ii) mediates aerobic respiration of atmospheric CO, and (iii) enhances survival of carbon-starved cells. On this basis, we confirm that atmospheric CO supports microbial survival and, with support from genomic, metagenomic, and metatranscriptomic analyses, propose a survival-centric model for the evolution and ecology of carboxydovores.

## Materials and Methods

### Bacterial strains and growth conditions

**Table S7** lists the bacterial strains and plasmids used in this study. *Mycobacterium smegmatis* mc^2^155 [66] and the derived strain Δ*coxL* were maintained on lysogeny broth (LB) agar plates supplemented with 0.05% (w/v) Tween80. For broth culture, *M. smegmatis* was grown on Hartmans de Bont minimal medium [67] supplemented with 0.05% (w/v) tyloxapol and 5.8 mM glycerol. *Escherichia coli* TOP10 cells were maintained on LB agar plates and grown in LB broth. Liquid cultures of both *M. smegmatis* and *E. coli* were incubated on a rotary shaker at 200 rpm, 37°C unless otherwise specified. Selective LB or LBT media used for cloning experiments contained gentamycin at 5 μg mL^−1^ for *M. smegmatis* and 20 μg mL^−1^ for *E. coli*.

### Mutant construction

A markerless deletion of the *coxL* gene (MSMEG_0746) was constructed by allelic exchange mutagenesis. Briefly, a 2245 bp fragment containing the fused left and right flanks of the MSMEG_0746 gene was synthesized by GenScript. This fragment was cloned into the SpeI site of the mycobacterial shuttle plasmid pX33 [68] with *E. coli* TOP10 and transformed into *M. smegmatis* mc^2^155 electrocompetent cells. To allow for temperature-sensitive vector replication, the transformants were incubated on LBT-gentamycin agar at 28°C for five days until colonies were visible. Catechol-reactive colonies were sub-cultured on to LBT-gentamycin agar plates incubated at 40°C for three days to facilitate the first recombination of the *coxL* flanks into the chromosome. To allow the second recombination and removal of the backbone vector to occur, colonies that were gentamycin-resistant and catechol-reactive were sub-cultured in LBT-sucrose agar and incubated at 40°C for three days. The resultant colonies were screened by PCR to discriminate Δ*coxL* mutants from wild-type revertants **(Figure S1)**. Whole-genome sequencing (Peter Doherty Institute, University of Melbourne) confirmed *coxL* was deleted and no other SNPs were present in the Δ*coxL* strain. **Table S8** lists the cloning and screening primers used in this study.

### Shotgun proteome analysis

For shotgun proteome analysis, 500 mL cultures of *M. smegmatis* were grown in triplicate in 2.5 L aerated conical flasks. Cells were harvested at mid-exponential phase (OD_600_ ~ 0.25) and mid-stationary phase (72 hours post OD_max_ ~0.9) by centrifugation (10,000 × *g*, 10 min, 4°C). They were subsequently washed in phosphate-buffered saline (PBS; 137 mM NaCl, 2.7 mM KCl, 10 mM Na_2_HPO_4_ and 2 mM KH_2_PO_4_, pH 7.4), recentrifuged, and resuspended in 8 mL lysis buffer (50 mM Tris-HCl, pH 8.0, 1 mM PMSF, 2 mM MgCl_2_, 5 mg mL^−1^ lysozyme, 1 mg DNase). The resultant suspension was then lysed by passage through a Constant Systems cell disruptor (40,000 psi, four times), with unbroken cells removed by centrifugation (10,000 × *g*, 20 min, 4°C). To denature proteins, lysates were supplemented with 20% SDS to a final concentration of 4%, boiled at 95°C for 10 min, and sonicated in a Bioruptor (Diagenode) using 20 cycles of ‘30 seconds on’ followed by ‘30 seconds off’. The lysates were clarified by centrifugation (14,000 × *g*, 10 mins). Protein concentration was confirmed using the bicinchoninic acid assay kit (Thermo Fisher Scientific) and equal amounts of protein were processed from both exponential and stationary phase samples for downstream analyses. After removal of SDS by chloroform/methanol precipitation, the proteins were proteolytically digested with trypsin (Promega) and purified using OMIX C18 Mini-Bed tips (Agilent Technologies) prior to LC-MS/MS analysis. Using a Dionex UltiMate 3000 RSL Cnano system equipped with a Dionex UltiMate 3000 RS autosampler, the samples were loaded via an Acclaim PepMap 100 trap column (100 μm × 2 cm, nanoViper, C18, 5 μm, 100 Å; Thermo Scientific) onto an Acclaim PepMap RSLC analytical column (75 μm × 50 cm, nanoViper, C18, 2 μm, 100 Å; Thermo Scientific). The peptides were separated by increasing concentrations of buffer B (80% acetonitrile / 0.1% formic acid) for 158 min and analyzed with an Orbitrap Fusion Tribrid mass spectrometer (Thermo Scientific) operated in data-dependent acquisition mode using in-house, LFQ-optimized parameters. Acquired.raw files were analyzed with MaxQuant [69] to globally identify and quantify proteins across the two conditions. Data visualization and statistical analyses were performed in Perseus [70].

### Activity staining

For CO dehydrogenase activity staining, 500 mL cultures of wild-type and Δ*coxL M. smegmatis* were grown to mid-stationary phase (72 hours post OD_max_ ~0.9) in 2.5 L aerated conical flasks. Cells were harvested by centrifugation, resuspended in lysis buffer, and lysed with a cell disruptor as described above. Following removal of unlysed cells by centrifugation (10,000 × *g*, 20 min, 4°C), the whole-cell lysates were fractionated into cytosols and membranes by ultracentrifugation (150,000 × *g*). The protein concentration of the lysates, cytosols, and membranes was determined using the bicinchoninic acid assay [71] against bovine serum albumin standards. Next, 20 μg protein from each fraction was loaded onto native Bis-Tris polyacrylamide gels (7.5% w/v running gel, 3.75% w/v stacking gel) prepared as described elsewhere [72] and run alongside a protein standard (NativeMark Unstained Protein Standard, Thermo Fisher Scientific) at 25 mA for 3 hr. For total protein staining, gels were incubated in AcquaStain Protein Gel Stain (Bulldog Bio) at 4°C for 3 hr. For CO dehydrogenase staining [14], gels were incubated in 50 mM Tris-HCl buffer containing 50 μM nitroblue tetrazolium chloride (NBT) and 100 μM phenazine methosulfate in an anaerobic jar (100% CO v/v atmosphere) at room temperature for 24 hours. Weak bands corresponding to CO dehydrogenase activity were also observed for wild-type fractions after 4 hours.

### Gas chromatography

Gas chromatography was used to determine the kinetics and threshold of CO dehydrogenase activity of *M. smegmatis*. Briefly, 30 mL stationary-phase cultures of wild-type and Δ*coxL M. smegmatis* strains were grown in 120 mL serum vials sealed with butyl rubber stoppers. At 72 hours post-OD_max_, cultures were reaerated (1 h), resealed, and amended with CO (*via* 1% v/v CO in N_2_ gas cylinder, 99.999% pure) to achieve headspace concentrations of ~200 ppmv. Cultures were agitated (150 rpm) for the duration of the incubation period to enhance CO transfer to the cultures and maintain an aerobic environment. Headspace samples of 1 mL were periodically collected using a gas-tight syringe to measure CO. Gas concentrations in samples were measured by gas chromatography using a pulsed discharge helium ionization detector (model TGA-6791-W-4U-2, Valco Instruments Company Inc.) as previously described [44]. Concentrations of CO in each sample were regularly calibrated against ultra-pure CO gas standards of known concentrations to the limit of detection of 9 ppbv CO. Kinetic analysis was performed as described, except cultures were amended with six different starting concentrations of CO (4000, 2000, 1000, 500, 200, 50 ppmv) and oxidation was measured at up to five timepoints (0, 2, 4, 6, 8 h). Reaction velocity relative to the gas concentration was calculated at each timepoint and plotted on a Michaelis-Menten curve. *V*_max app_ and *K*_m app_ values were derived through a non-linear regression model (GraphPad Prism, Michaelis-Menten, least squares fit) and linear regressions based on Lineweaver-Burk, Eadie-Hofstee, and Hanes-Woolf plots.

### Respirometry measurements

For respirometry measurements, 30 mL cultures of wild-type and Δ*coxL M. smegmatis* were grown to mid-stationary phase (72 hours post OD_max_ ~0.9) in 125 mL aerated conical flasks. Rates of O_2_ consumption were measured before and after CO addition using a Unisense O_2_ microsensor. Prior to measurement, the electrode was polarized at −800 mV for 1 hour with a Unisense multimeter and calibrated with O_2_ standards of known concentration. Gas-saturated PBS was prepared by bubbling PBS with 100% (v/v) of either O_2_ or CO for 5 min. Initially, O_2_ consumption was measured in 1.1 mL microrespiration assay chambers sequentially amended with *M. smegmatis* cell suspensions (0.9 mL) and O_2_-saturated PBS (0.1 mL) that were stirred at 250 rpm at room temperature. After initial measurements, 0.1 mL of CO-saturated PBS was added into the assay mixture. Changes in O_2_ concentrations were recorded using Unisense Logger Software (Unisense, Denmark). Upon observing a linear change in O_2_ concentration, rates of consumption were calculated over a period of 20 s and normalized against total protein concentration.

### Gene expression analysis

To assess CO dehydrogenase gene expression by qRT-PCR, synchronized 30 mL cultures of *M.* smegmatis were grown in triplicate in either 125 mL aerated conical flasks or 120 mL sealed serum vials supplemented with 1% (w/v) CO. Cultures were quenched at mid-exponential phase (OD_600_ ~0.25) or mid-stationary phase (three days post-OD_max_ ~0.9) with 60 mL cold 3:2 glycerol:saline solution (−20°C). They were subsequently harvested by centrifugation (20,000 × *g*, 30 minutes, −9°C), resuspended in 1 mL cold 1:1 glycerol:saline solution (−20°C), and further centrifuged (20,000 × *g*, 30 minutes, −9°C). For cell lysis, pellets were resuspended in 1 mL TRIzol Reagent, mixed with 0.1 mm zircon beads, and subjected to five cycles of bead-beating (4,000 rpm, 30 seconds) in a Biospec Mini-Beadbeater. Total RNA was subsequently extracted by phenol-chloroform extraction as per manufacturer’s instructions (TRIzol Reagent User Guide, Thermo Fisher Scientific) and resuspended in diethylpyrocarbonate (DEPC)-treated water. RNA was treated with DNase using the TURBO DNA-free kit (Thermo Fisher Scientific) as per the manufacturer’s instruction.

RNA concentration, purity, and integrity were confirmed by using a NanoDrop ND-1000 spectrophotometer and running extracts on a 1.2% agarose gel. cDNA was then synthesized using SuperScript III First-Strand Synthesis System for qRT-PCR (Thermo Fisher Scientific) with random hexamer primers as per the manufacturer’s instructions. qPCR was used to quantify the copy numbers of the target gene *coxL* and housekeeping gene *sigA* against amplicon standards of known concentration. A standard curve was created based on the cycle threshold (Ct) values of *coxL* and *sigA* amplicons that were serially diluted from 10^8^ to 10 copies (R^2^ > 0.99). The copy number of the genes in each sample was interpolated based on each standard curve and values were normalized to *sigA* expression in exponential phase in ambient air. For each biological replicate, all samples, standards, and negative controls were run in technical duplicate. All reactions were run in a single 96-well plate using the PowerUp SYBR Green Master Mix (Thermo Fisher Scientific) and LightCycler 480 Instrument (Roche) according to each manufacturers’ instructions.

### Growth and survival assays

For growth and survival assays, cultures were grown in 30 mL media in either 125 mL aerated conical flasks or 120 mL sealed serum vials containing an ambient air headspace amended with 20% (v/v) CO. Growth was monitored by measuring optical density at 600 nm (1 cm cuvettes; Eppendorf BioSpectrometer Basic); when OD_600_ was above 0.5, cultures were diluted ten-fold in before measurement. All growth experiments were performed using three biological replicates. To count colony forming units (CFU mL^−1^), each culture was serially diluted in HdB (no carbon source) and spotted on to agar plates in technical quadruplicates. Survival experiments were performed on two separate occasions using three biological replicates in the first experiment and six biological replicates in the second experiment. Percentage survival was calculated for each replicate by dividing the CFU mL^−1^ at each timepoint with the CFU mL^−1^ count at OD_max_.

### Glycerol quantification

Glycerol concentration in media was measured colorimetrically. Samples of 900 μL were taken periodically from triplicate cultures during growth, cells were pelleted (9,500 x *g*, 2 minutes) and supernatant was collected and stored at −20°C. Glycerol content for all supernatant samples was measured simultaneously in a single 96-well plate using a Glycerol Assay Kit (Sigma-Aldrich) as per manufacturer’s instructions. Absorbance was measured at 570 nm using an Epoch 2 microplate reader (BioTek). A standard curve was constructed using four standards of glycerol (0 mM, 0.3 mM, 0.6 mM and 1 mM; R^2^ > 0.99). Glycerol concentration was interpolated from this curve. Samples were diluted either five-fold or two-fold in UltraPure water such that they fell within the curve. All samples, standards and blanks were run in technical duplicate.

### Genome survey

The amino acid sequences of the catalytic subunits of all putative form I CO dehydrogenases (CoxL) represented in the National Center for Biotechnology Information (NCBI) Reference Sequence (RefSeq) [73]. All sequences with greater than 55% sequence identity and 90% query coverage to CoxL sequences of *Oligotropha carboxidovorans* (WP_013913730.1), *Mycobacterum smegmatis* (WP_003892166.1), and *Natronorubrum bangense* (WP_006067999.1) were retrieved by protein BLAST [74]. Homologous sequences with less than 55% sequence encoded form II CO dehydrogenases and hence were not retrieved. The dataset was manually curated to dereplicate sequences within species and remove incomplete sequences. The final dataset contained a total of 709 CoxL sequences across 685 different bacterial and archaeal species **(Table S3)**.

### Phylogenetic analysis

To construct phylogenetic trees, the retrieved sequences were aligned using ClustalW in MEGA7 [75]. Initially, the phylogenetic relationships of 709 sequences were visualized on a neighbor-joining tree based on the Poisson correction method and bootstrapped with 500 replicates. Subsequently, the phylogenetic relationships of a representative subset of 94 sequences were visualized on a maximum-likelihood tree based on the Poisson correction method and bootstrapped with 200 replicates. Both trees were rooted with the protein sequences of five form II CO dehydrogenase catalytic subunit sequences (WP_012893108.1, WP_012950878.1, WP_013076571.1, WP_01359081.1, WP_013388721.1). We confirmed that trees of similar topology were produced upon using a range of phylogenetic methods, namely neighbor-joining, maximum-parsimony and maximum-likelihood in MEGA, Mr Bayes, phyml, and iqtree. In addition, equivalent trees were created by using the protein sequences of the CO dehydrogenase medium subunit (CoxM), small subunits (CoxS), or concatenations of all three subunits (CoxLMS). Varying the form II CO dehydrogenase sequence used also had no effect on the overall topology.

### Metagenome and metatranscriptome analysis

Forty pa-irs of metagenomes and metatranscriptomes that encompassed a range of soil and marine sample types were selected and downloaded from the Joint Genome Institute (JGI) Integrated Microbial Genomes System [76] and the NCBI Sequence Read Archive (SRA) [77]. **Table S5** provides details of the datasets used. Raw metagenomes and metatranscriptomes were subjected to quality filtering using NGS QC Toolkit [78] (version 2.3.3, default settings, i.e. base quality score and read length threshold are 20 and 70%, respectively). SortMeRNA [79] (version 2.1, default settings and default rRNA databases) was used to removed ribosomal RNA (rRNA) reads from metatranscriptomes. Each metagenome and metatranscriptome was subsampled to an equal depth of 5 million reads and 2 million reads, respectively, using seqtk (https://github.com/lh3/seqtk) seeded with parameter -s100. Subsampled datasets were then screened in DIAMOND (version 0.9.24.125, default settings, one maximum target sequence per query) [80] using the 709 CoxL protein sequences **(Table S3)** and the 3261 hydrogenase catalytic subunit gene sequences from HydDB [81]. Hits to CoxL were filtered with an amino acid alignment length over 40 residues and a sequence identity over 60%. Clade classification of the reads was based on their closest match to the CoxL sequence dataset. Hydrogenase hits were filtered with the same amino acid alignment length cutoff and a sequence identity over 50%. Group 4 [NiFe]-hydrogenase hits with a sequence identity below 60% were discarded.

## Results

### *Mycobacterium smegmatis* synthesizes carbon monoxide dehydrogenase during a coordinated response to organic carbon starvation

We first performed a proteome analysis to gain a system-wide context of the levels of CO dehydrogenase during growth and survival of *M. smegmatis*. Shotgun proteomes were compared for triplicate cultures grown in glycerol-supplemented minimal media under two conditions: mid-exponential growth (OD_600_ ~ 0.25; 5.1 mM glycerol left in medium) and mid-stationary phase following carbon limitation (72 hours post OD_max_ ~0.9; no glycerol detectable in medium) (Fig. 1a). There was a major change in the proteome profile, with 270 proteins more abundant and 357 proteins less abundant by at least four-fold (*p* < 0.05) in the carbon-limited condition (Fig. 1b; **Table S2**).

**Figure 1.**
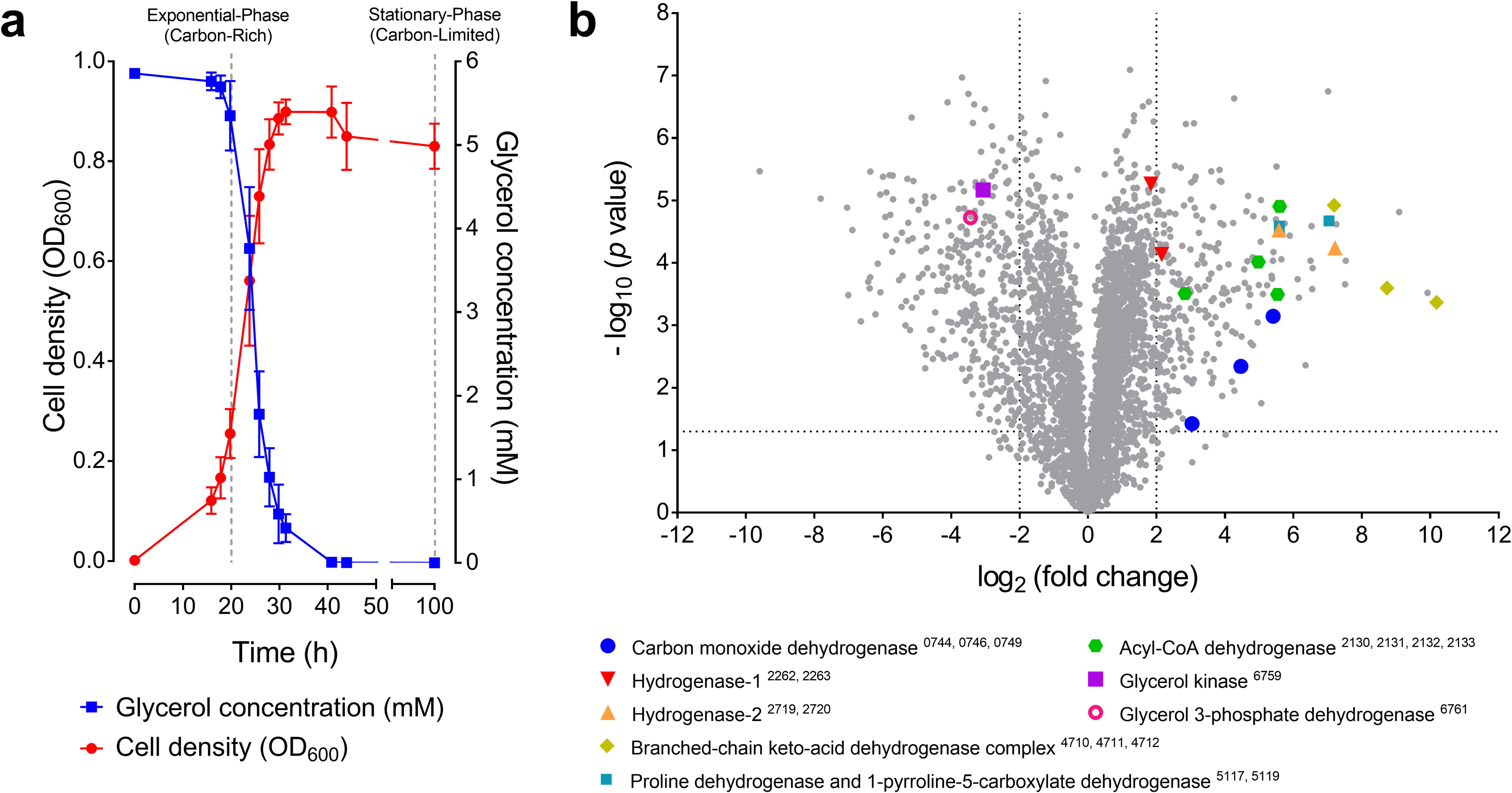
Comparison of proteome composition of carbon-replete and carbon-limited cultures of *Mycobacterium smegmatis*. (**a**) Growth of *M. smegmatis* in Hartmans de Bont minimal medium supplemented with 5.8 mM glycerol. The glycerol concentration of the external medium is shown. Error bars show standard deviations of three biological replicates. Cells were harvested for proteomic analysis at OD_600_ = 0.25 (mid-exponential phase, glycerol-rich) and three days post OD_max_ (mid-stationary phase, glycerol-limited). (**b**) Volcano plot showing relative expression change of genes following carbon-limitation. Fold change was determined by dividing the relative abundance of each protein in three stationary phase proteomes with that in the three exponential phase proteomes (biological replicates). Each protein is represented by a grey dot. Structural subunits of selected metabolic enzymes, including the form I CO dehydrogenase, are highlighted and their locus numbers are shown in subscript in the legend.

The top 50 proteins with increased abundance included those involved in trace gas metabolism and amino acid catabolism. In line with our hypotheses, there was an increase in the structural subunits encoding a putative form I CO dehydrogenase, including a 54-fold increase in the catalytic subunit CoxL. Levels of the two uptake hydrogenases also increased, particularly the catalytic subunit of hydrogenase-2 (HhyL, 148-fold), in line with previous observations that mycobacteria persist on atmospheric H_2_ [54, 63]. There was also evidence that *M. smegmatis* generates additional reductant in this condition by catabolizing amino acid reserves: the three subunits of a branched-chain keto-acid dehydrogenase complex were the most differentially abundant proteins overall and there was also a strong induction of the proline degradation pathway, including the respiratory proline dehydrogenase (Fig. 1b).

The abundance of various enzymes mediating organic carbon catabolism decreased, including the respiratory glycerol 3-phosphate dehydrogenase (10-fold) and glycerol kinase (8-fold), in line with cultures having exhausted glycerol supplies (Fig. 1b). The proteome also suggests that various energetically-expensive processes, such as cell wall, ribosome, and DNA synthesis, were downregulated **(Table S2)**. Overall, these results suggest that *M. smegmatis* reduces its energy expenditure and expands its metabolic repertoire, including by oxidizing CO, to stay energized during starvation.

### Carbon monoxide dehydrogenase mediates atmospheric CO oxidation and supports aerobic respiration

Having confirmed that a putative CO dehydrogenase is present in stationary-phase *M. smegmatis* cells, we subsequently confirmed its activity through whole-cell biochemical assays. To do so, we constructed a markerless deletion of the *coxL* gene (MSMEG_0746) **(Fig. S1)**. Native polyacrylamide gels containing fractions of wild-type *M. smegmatis* harvested in carbon-limited stationary-phase cells strongly stained for CO dehydrogenase activity in a 100% CO atmosphere; the molecular weight of the band corresponds to the theoretical molecular weight of a dimer of CoxLMS subunits (~269 kDa). However, no activity was observed in the Δ*coxL* background (Fig. 2a).

**Figure 2.**
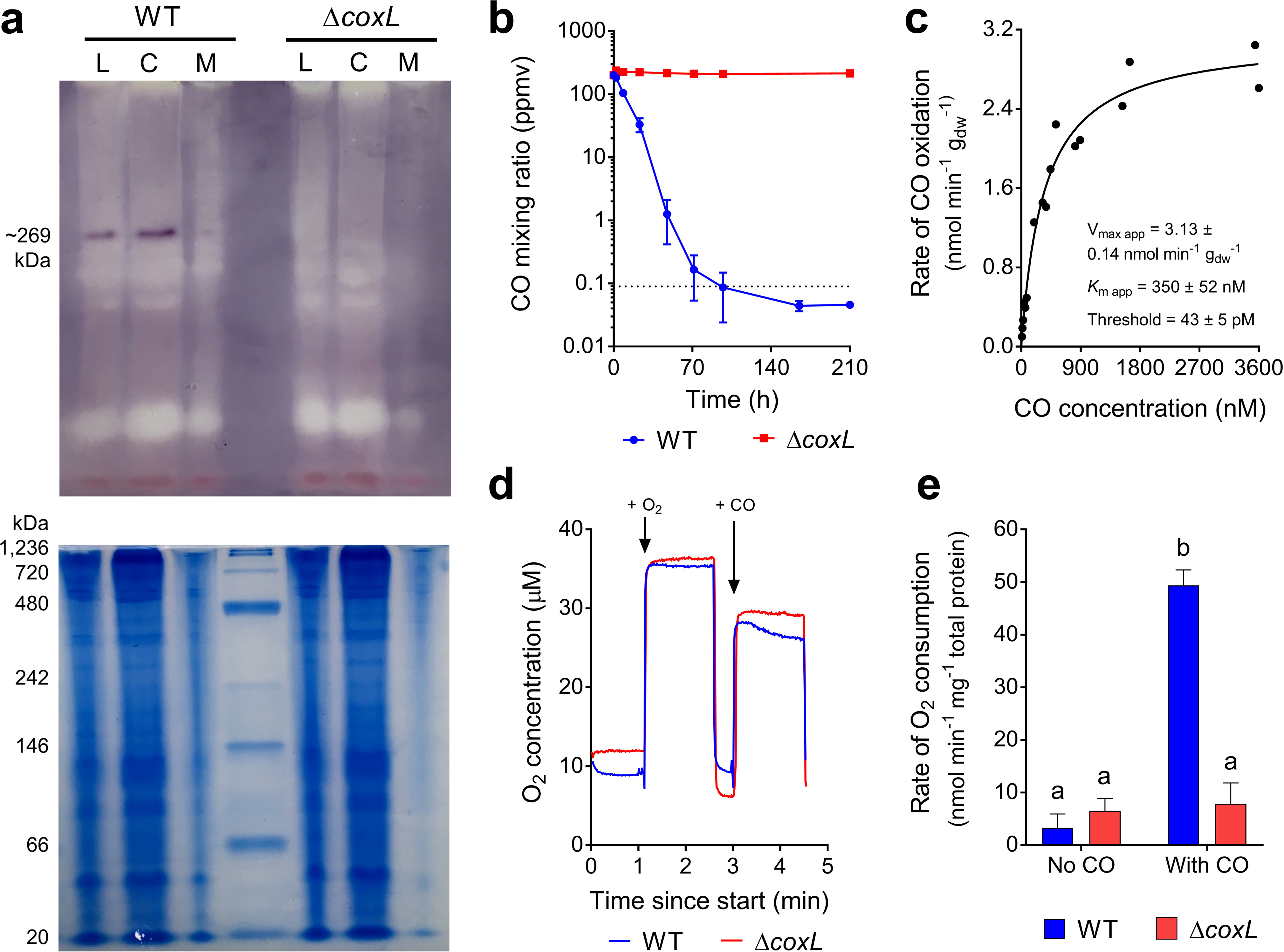
Comparison of carbon monoxide dehydrogenase activity of *Mycobacterium smegmatis* wild-type and Δ*coxL* cultures. (**a**) Zymographic observation of CO dehydrogenase activity and localization. The upper gel shows enzyme activity stained with the artificial electron acceptor nitroblue in a CO-rich atmosphere. The lower gel shows protein ladder and whole protein stained with Coomassie Blue. Results are shown for whole-cell lysates (L), cytosolic fractions (C), and membrane fractions (M) of wild-type (WT) and Δ*coxL* cultures. (**b**) Gas chromatography measurement of CO oxidation to sub-atmospheric levels. Mixing ratios are displayed on a logarithmic scale, the dotted line shows the average atmospheric mixing ratios of CO (90 ppbv), and error bars show standard deviations of three biological replicates. (**c**) Apparent kinetic parameters of CO oxidation by wild-type cultures. Curves of best fit and kinetic parameters were calculated based on a Michaelis-Menten non-linear regression model. *V*_max app_ and *K*_m app_ values derived from other models are shown in **Table S4**. (**d**) Examples of traces from oxygen electrode measurements. O_2_ levels were measured before and after CO addition in both a wild-type and Δ*coxL* background. (**e**) Summary of rates of O_2_ consumption measured using an oxygen electrode. Centre values show means and error bars show standard deviations from three biological replicates. For all values with different letters, the difference between means is statistically significant (*p* < 0.001) based on student’s t-tests.

Gas chromatography measurements confirmed that *M. smegmatis* oxidized carbon monoxide at atmospheric concentrations. Stationary-phase cultures oxidized the CO supplemented in the headspace (~200 ppmv) to sub-atmospheric concentrations (46 ± 5 ppbv) within 100 hours (Fig. 2b). The apparent kinetic parameters of this activity (*V*_max app_ = 3.13 nmol g_dw_^−1^ min^−1^; *K*_m app_ = 350 nM; threshold _app_ = 43 pM) are consistent with a moderate-affinity, slow-acting enzyme (Fig. 2c; **Table S4**). The rates are similar to those previously measured for hydrogenase-2 [63]. No change in CO mixing ratios was observed for the Δ*coxL* strain (Fig. 2b), confirming that the form I CO dehydrogenase is the sole CO-oxidizing enzyme in *M. smegmatis*. In turn, these results provide the first genetic proof that form I CO dehydrogenases mediate atmospheric CO oxidation.

We performed oxygen electrode experiments to confirm whether CO addition stimulated aerobic respiration. In stationary-phase cultures, addition of CO caused a 15-fold stimulation of respiratory O_2_ consumption relative to background rates (*p* < 0.0001). This stimulation was observed in the wild-type strain, but not the Δ*coxL* mutant, showing it is dependent on CO oxidation activity of the CO dehydrogenase (Fig. 2d & 2e). Thus, while this enzyme is predominantly localized in the cytosol (Fig. 2a), it serves as a *bona fide* respiratory dehydrogenase that supports aerobic respiration in *M. smegmatis*.

### Carbon monoxide is dispensable for growth and detoxification, but enhances survival during carbon starvation

We then performed a series of experiments to resolve the expression and importance of the CO dehydrogenase during growth and survival. Consistent with the proteomic analyses, expression levels of *coxL* were low in carbon-replete cultures (mid-exponential phase; 1.35 × 10^7^ transcripts g_dw_^−1^) and increased 56-fold in carbon-limited cultures (mid-stationary phase; 7.48 × 10^8^ transcripts g_dw_^−1^; *p* < 0.01). Addition of 1% CO did not significantly change *coxL* expression in either growing or stationary cultures (Fig. 3a). These profiles suggest that *M. smegmatis* expresses CO dehydrogenase primarily to enhance survival by scavenging atmospheric CO, rather than to support growth on elevated levels of CO.

**Figure 3.**
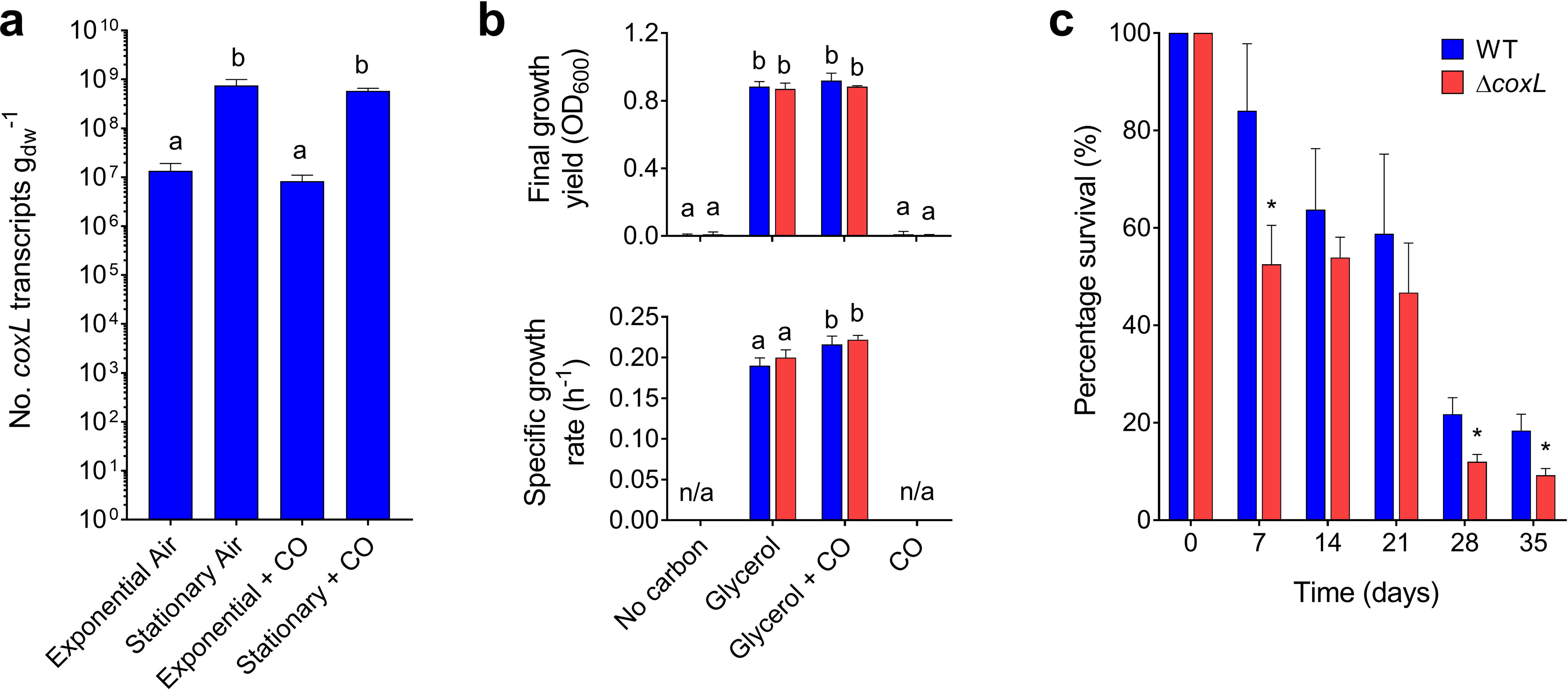
Expression and importance of carbon monoxide dehydrogenase during growth and survival of *Mycobacterium smegmatis*. (**a**) Normalized number of transcripts of the CO dehydrogenase large subunit gene (*coxL*; MSMEG_0746) in wild-type cultures harvested during exponential phase (carbon-replete) and stationary phase (carbon-limited) in the presence of either ambient CO or 1% CO. Error bars show standard deviations of four biological replicates. For all values with different letters, the difference between means is statistically significant (*p* < 0.01) based on student’s t-tests. (**b**) Final growth yields (OD_max_) and specific growth rates wild-type and Δ*coxL* strains. Strains were grown on Hartman de Bont minimal medium supplemented with either 5.5 mM glycerol, 20% CO, or both 5.5 mM glycerol and 20% CO. Values labelled with different letters are significantly different (*p* < 0.05) based on student’s t-tests. Error bars show standard deviations of three biological replicates. (**c**) Long-term survival of wild-type and Δ*coxL* strains in Hartman de Bont minimal medium supplemented with either 5.5 mM glycerol. Percentage survival was calculated by dividing the colony forming units (CFU mL^−1^) at each timepoint with those counted at OD_max_ (day 0). Error bars show standard deviations of nine biological replicates. For asterisked values, there was a significant difference in survival of Δ*coxL* strains compared to the wild-type (*p* < 0.05) based on student’s t-tests.

These inferences were confirmed by monitoring the growth of the wild-type and Δ*coxL* strains under different conditions. The strains grew identically on glycerol-supplemented minimal medium. Addition of 20% CO caused a slight increase in doubling time for both strains and did not affect growth yield (Fig. 3b). This suggests that *M. smegmatis* is highly tolerant of CO but does not require CO dehydrogenase to detoxify it. *M. smegmatis* did not grow chemolithoautotrophically on a minimal medium with 20% CO as the sole carbon and energy source (Fig. 3b). While carboxydotrophic growth was previously reported for this strain, the authors potentially observed CO-tolerant heterotrophic or mixotrophic growth, given the reported media contained metabolizable organic carbon sources [40]. Consistently, *M. smegmatis* lacks key enzymes of the Calvin-Benson cycle (e.g. RuBisCO, ribulose 1,5-bisphosphate carboxylase) typically required for carboxydotrophic growth.

Finally, we monitored the long-term survival of the two strains after they reached maximum cell counts upon exhausting glycerol supplies (Fig. 1a). The percentage survival of the Δ*coxL* strain was lower than the wild-type at all timepoints, including by 45% after four weeks and 50% after five weeks of persistence. These findings were reproducible across two independent experiments and were significant at the 98% confidence level (Fig. 3c). Such reductions in relative percentage survival are similar to those previously observed for uptake hydrogenase mutants in *M. smegmatis* (47%) [53, 54] and *Streptomyces avermilitis* (74%) [57]. These experiments therefore provide genetic proof that atmospheric CO oxidation mediated by form I CO dehydrogenases enhances bacterial persistence.

### Atmospheric carbon monoxide oxidation is an ancient, taxonomically widespread and ecologically important process

We subsequently surveyed genomic, metagenomic, and metatranscriptomic datasets to gain insights the taxonomic and ecological distribution of atmospheric CO oxidation. This yielded 709 amino acid sequences encoding large subunits of the form I CO dehydrogenases (CoxL) across some 685 species, 196 genera, 49 orders, and 25 classes of bacteria and archaea (**Table S3;** Fig. 4a & 4b). The retrieved sequences encompassed all sequenced species, across seven phyla (Figure 4b) that have previously been shown to mediate aerobic CO oxidation **(Table S1)**. We also detected *coxL* genes in nine other phyla where aerobic CO oxidation has yet to be experimentally demonstrated (Fig. 4b). Hence, the capacity for aerobic CO respiration appears to be a much more widespread trait among aerobic bacteria and archaea than previously reported [49, 62]. It is particularly notable that *coxL* genes were detected in representatives of seven of the nine [64, 82] most dominant soil phyla, namely Proteobacteria, Actinobacteriota, Acidobacteriota, Chloroflexota, Firmicutes, Gemmatimonadota, and Bacteroidota (Fig. 4b).

**Figure 4.**
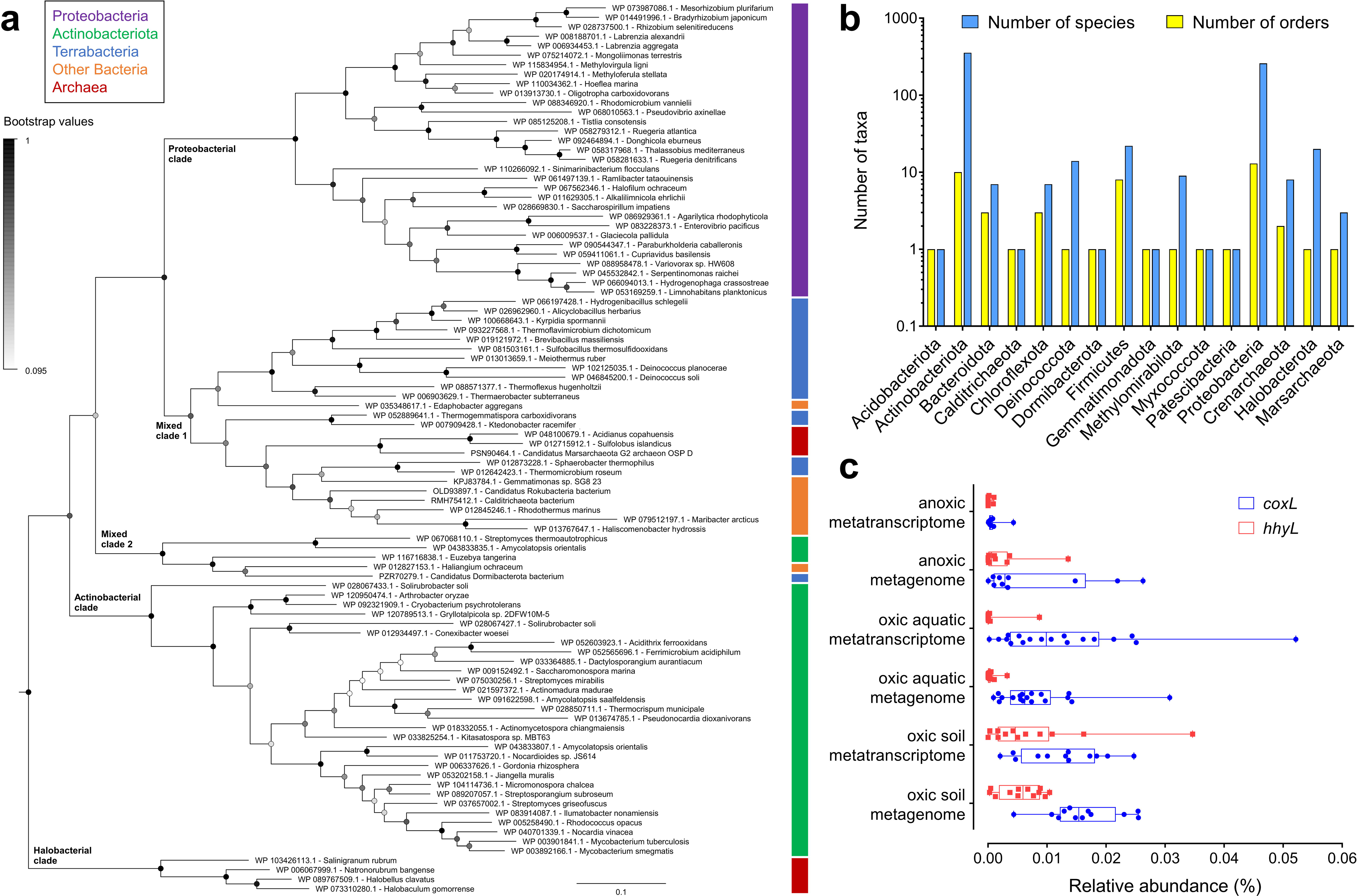
Distribution of carbon monoxide dehydrogenases in genomes, metagenomes, and metatranscriptomes. (**a**) Maximum-likelihood phylogenetic tree showing the evolutionary history of the catalytic subunit of the form I CO dehydrogenase (CoxL). Evolutionary distances were computed using the Poisson correction model, gaps were treated by partial deletion, and the tree was bootstrapped with 200 replicates. The tree was constructed using a representative subset of 94 CoxL amino acid sequences from **Table S3** and a neighbor-joining tree containing all 709 CoxL sequences retrieved in this study is provided in **Fig. S2**. The major clades of the tree are labeled, and the colored bars represent the phylum that each sequence is affiliated with. The tree was rooted with five form II CO dehydrogenase sequences (not shown). (**b**) Phylum-level distribution of the CoxL-encoding species and orders identified in this work. (**c**) Abundance of *coxL* genes and transcripts in environmental samples. In total, 40 pairs of metagenomes and metatranscriptomes (20 aquatic, 20 terrestrial) were analyzed from a wide range of biomes (detailed in **Table S5**). The abundance of *hhyL* genes and transcripts, encoding the high-affinity group 1h [NiFe]-hydrogenase, are shown for comparison. Box plots show the individual values and their mean, quartiles, and range for each dataset.

We constructed phylogenetic trees to visualize the evolutionary relationships of CoxL protein sequences (Fig. 4a; **Fig. S2**). The trees contained five monophyletic clades that differed in phylum-level composition, namely actinobacterial, proteobacterial, and halobacterial clades, as well as mid-branching major (mixed 1) and minor (mixed 2) clades of mixed composition containing representatives from seven and three different phyla respectively. Clades were well-supported by bootstrap values, with exception of the mixed 2 clade (Fig. 4a; **Fig. S2)**. Trees with equivalent clades were produced when using seven distinct phylogenetic methods, using other CO dehydrogenase subunits (CoxM, CoxS, and CoxLMS concatenations), or varying the outgroup sequences. In all cases, major clades included CoxL proteins of at least one previously characterized carboxydotroph or carboxydovore **(Table S1)**. Surprisingly, all clades also contained species that have been previously shown to oxidize atmospheric CO **(Table S1)**. This suggests that atmospheric CO oxidation is a widespread and ancestral capability among CO dehydrogenases. In contrast, CO dehydrogenases known to support aerobic carboxydotrophic growth were sparsely distributed across the tree (Fig. 4a; **Table S1**).

To better understand the ecological significance of aerobic CO oxidation, we surveyed the abundance of *coxL* sequences across 40 pairs of metagenomes and metatranscriptomes **(Table S5)**. Genes and transcripts for *coxL* were detected across a wide range of biomes. They were particularly abundant in the oxic terrestrial and marine samples surveyed (1 in every 8,000 reads), for example grassland and rainforest soils, coastal and mesopelagic seawater, and salt marshes **(Table S6)**. In contrast, they were expressed at very low levels in anaerobic samples (e.g. groundwater, deep subsurface, peatland) **(Figure S3)**. Across all surveyed metatranscriptomes, the majority of the *coxL* hits were affiliated with the mixed 1 (40%), proteobacterial (25%), and actinobacterial (25%) clades, with minor representation of the mixed 2 (8%) and halobacterial (2%) clades **(Table S6)**. The normalized transcript abundance of *coxL* was higher than the genetic determinants of atmospheric H2 oxidation (*hhyL*; high-affinity hydrogenase) in most samples (18-fold in aquatic samples, 1.2-fold in terrestrial samples) (Fig. 4c). Together, this suggests that CO oxidation is of major importance in aerated environments and is mediated by a wide range of bacteria and archaea.

## Discussion

In this work, we validated that atmospheric CO oxidation supports bacterial survival during nutrient limitation. *M. smegmatis* increases the transcription and synthesis of a form I CO dehydrogenase by 50-fold as part of a coordinated response to organic carbon limitation. Biochemical studies confirmed that this enzyme is kinetically adapted to scavenge atmospheric concentrations of CO and use the derived electrons to support aerobic respiration. In turn, genetic deletion of the enzyme did not affect growth under a range of conditions, but resulted in severe survival defects in carbon-exhausted cultures. These observations are reminiscent of previous observations that *M. smegmatis* expresses two high-affinity hydrogenases to persist by scavenging atmospheric H_2_ [53–55, 63]. In common with atmospheric H_2_, atmospheric CO is a high-energy, diffusible, and ubiquitous trace gas [28], and is therefore a dependable source of energy to sustain the maintenance needs of bacteria during persistence. Overall, the proteome results suggest that *M. smegmatis* activates CO scavenging as a core part of a wider response to enhance its metabolic repertoire; the organism appears to switch from acquiring energy organotrophically during growth to mixotrophically during survival by scavenging a combination of inorganic and organic energy sources.

In turn, it is probable that CO supports the persistence of many other bacterial and archaeal species. Atmospheric CO oxidation is a common trait among all carboxydovores tested to date and has been experimentally demonstrated in 18 diverse genera of bacteria and archaea [19, 29, 33, 36, 43, 44, 48]. In this regard, a recent study demonstrated that the hot spring bacterium *Thermomicrobium roseum* (phylum Chloroflexota) upregulates a form I CO dehydrogenase and oxidizes atmospheric CO as part of a similar response to carbon starvation [44]. It has also been demonstrated that the form I CO dehydrogenases of the known atmospheric CO scavenger *Ruegeria pomolori* [58] and a *Phaeobacter* isolate [59] from the marine *Roseobacter* clade (phylum Proteobacteria) are also highly upregulated under energy-limiting conditions. The capacity for atmospheric CO uptake has also been demonstrated in four halophilic archaeal genera (phylum Halobacterota) [33, 48] and may also extend to thermophilic archaea (phylum Crenarchaeota) [46, 47]. Moreover, two cultured aerobic methanotrophs harbour the capacity for aerobic CO respiration [83, 84]. Our study, by showing through a molecular genetic approach that CO oxidation enhances survival, provides a physiological rationale for these observations.

These results also have broader implications for understanding the biogeochemical cycling and microbial biodiversity at the ecosystem level. It is well-established that soil bacteria are major net sinks for atmospheric CO and marine bacteria mitigate geochemical oceanic emissions of this gas [10]. This study, by confirming the enzymes responsible and demonstrating that their activities support bacterial persistence, has ramifications for modelling these biogeochemical processes. In turn, we propose that CO is an important energy source supporting the biodiversity and stability of aerobic heterotrophic communities in terrestrial and aquatic environments. The genomic survey supports this by demonstrating that form I CO dehydrogenases, most of which are predicted to support atmospheric CO oxidation, are encoded by 685 species and 16 phyla of bacteria and archaea. In turn, the metagenomic and metatranscriptomic analyses confirmed that *coxL* genes and transcripts are highly abundant in most aerated soil and marine ecosystems. The notably high abundance of *coxL* transcripts in pelagic samples of various depths suggests CO may be a major energy source for maintenance of marine bacteria. In soils, the oxidation of atmospheric CO may be of similar importance to atmospheric H_2_; this is suggested by the strength of the soil sinks for these gases [1, 85], the abundance of *coxL* and *hhyL* genes in soil metagenomes, and the distribution of these genes in the genomes of soil bacteria [86]. Atmospheric CO may be especially important for sustaining communities in highly oligotrophic soils, as indicated by previous studies in polar deserts [51], volcanic deposits [60, 62, 87], and salt flats [33, 88, 89]. Further work is now needed to understand which microorganisms mediate consumption of atmospheric CO *in situ* and how their activity is controlled by physicochemical factors.

Integrating these findings with the wider literature, we propose a new survival-centric model for the evolution of CO dehydrogenases. It was traditionally thought that aerobic CO oxidation primarily supports autotrophic and mixotrophic growth of microorganisms [11, 26]. However, the majority of studied CO-oxidizing bacteria are in fact carboxydovores, of which those that have been kinetically characterized can oxidize CO at sub-atmospheric levels **(Table S1)**. In turn, our phylogenomic analysis revealed that atmospheric CO-oxidizing bacteria are represented in all five clades of the phylogenetic tree, suggesting that the common ancestor of these enzymes also harbored sufficient substrate affinity to oxidize atmospheric CO. On this basis, we propose that microorganisms first evolved a sufficiently high-affinity form I CO dehydrogenase to subsist on low concentrations of CO. The genes encoding this enzyme were then horizontally and vertically disseminated to multiple bacterial and archaeal genera inhabiting different environments. On multiple occasions, certain bacterial lineages evolved to support growth on CO in microenvironments where present at elevated concentrations. This would have required relatively straightforward evolutionary innovations, namely acquisition of Calvin-Benson cycle enzymes (e.g. RuBisCO) and their integration with CO dehydrogenase. The modulation of CO dehydrogenase kinetics was likely not a prerequisite, given these enzymes efficiently oxidize CO at a wide range of substrate concentrations [19, 44], but may have subsequently enhanced carboxydotrophic growth. These inferences differ from hydrogenases, where high-affinity, oxygen-tolerant enzymes appear to have evolved from low-affinity, oxygen-sensitive ones [86]. However, it is probable that the processes of atmospheric CO and H_2_ oxidation evolved due to similar physiological pressures and over similar evolutionary timescales.

## Footnotes

### Author contributions

C.G. conceived this study. C.G., P.R.F.C., K.B., P.M.L., G.M.K., R.B.S., and C.H. designed experiments and analyzed data. C.G. and P.R.F.C. supervised students. Different authors were responsible for proteomic analysis (C.H., R.B.S., P.R.F.C., C.G.), knockout construction (P.R.F.C.), activity measurements (K.B., P.R.F.C., Z.I.), respirometry analysis (P.R.F.C.), expression profiling (K.B., P.R.F.C.), growth analysis (K.B.), survival assays (K.B.), genome surveys (C.G.), phylogenetic analysis (G.M.K., C.G.), and meta-omic analysis (P.M.L., C.G.). C.G., P.R.F.C., and K.B. wrote and edited the paper with input from all authors.

## Acknowledgements

This work was supported by an ARC DECRA Fellowship (DE170100310; awarded to C.G.), an NHMRC New Investigator Grant (APP5191146; awarded to C.G.), an Australian Government Research Training Program Stipend Scholarship (awarded to K.B. and Z.I.), and Monash University Doctoral Scholarships (awarded to P.R.F.C. and P.M.L.). We thank Dr George Taiaroa and A/Prof Debbie Williamson for sequencing the mutants, Blair Ney and Thanavit Jirapanjawat for their technical assistance, and Dr Eleonora Chiri for critically reading the manuscript.

The authors declare no conflict of interest.

## References

1. Khalil MAK, Rasmussen RA. The global cycle of carbon monoxide: Trends and mass balance. Chemosphere 1990; 20: 227–242.

2. Novelli PC, Masarie KA, Lang PM. Distributions and recent changes of carbon monoxide in the lower troposphere. J Geophys Res Atmos 1998; 103: 19015–19033.

3. Chi X, Winderlich J, Mayer J-C, Panov A V, Heimann M, Birmili W, et al. Long-term measurements of aerosol and carbon monoxide at the ZOTTO tall tower to characterize polluted and pristine air in the Siberian taiga. Atmos Chem Phys 2013; 13: 12271–12298.

4. Petrenko V V, Martinerie P, Novelli P, Etheridge DM, Levin I, Wang Z, et al. A 60 yr record of atmospheric carbon monoxide reconstructed from Greenland firn air. Atmos Chem Phys 2013; 13: 7567–7585.

5. Bartholomew GW, Alexander M. Soil as a sink for atmospheric carbon monoxide. Science 1981; 212: 1389–1391.

6. Inman RE, Ingersoll RB, Levy EA. Soil: a natural sink for carbon monoxide. Science 1971; 172: 1229–1231.

7. Kirschke S, Bousquet P, Ciais P, Saunois M, Canadell JG, Dlugokencky EJ, et al. Three decades of global methane sources and sinks. Nat Geosci 2013; 6: 813–823.

8. Swinnerton JW, Linnenbom VJ, Lamontagne RA. The ocean: a natural source of carbon monoxide. Science 1970; 167: 984–986.

9. Xie H, Bélanger S, Demers S, Vincent WF, Papakyriakou TN. Photobiogeochemical cycling of carbon monoxide in the southeastern Beaufort Sea in spring and autumn. Limnol Oceanogr 2009; 54: 234–249.

10. Zafiriou OC, Andrews SS, Wang W. Concordant estimates of oceanic carbon monoxide source and sink processes in the Pacific yield a balanced global “blue-water” CO budget. Global Biogeochem Cycles 2003; 17.

11. King GM, Weber CF. Distribution, diversity and ecology of aerobic CO-oxidizing bacteria. Nat Rev Microbiol 2007; 5: 107–118.

12. Zavarzin GA, Nozhevnikova AN. Aerobic carboxydobacteria. Microb Ecol 1977; 3: 305–326.

13. Meyer O, Schlegel HG. Reisolation of the carbon monoxide utilizing hydrogen bacterium *Pseudomonas carboxydovorans* (Kistner) comb. nov. Arch Microbiol 1978; 118: 35–43.

14. Lorite MJ, Tachil J, Sanjuán J, Meyer O, Bedmar EJ. Carbon monoxide dehydrogenase activity in *Bradyrhizobium japonicum*. Appl Environ Microbiol 2000; 66: 1871–1876.

15. Kiessling M, Meyer O. Profitable oxidation of carbon monoxide or hydrogen during heterotrophic growth of *Pseudomonas carboxydoflava*. FEMS Microbiol Lett 1982; 13: 333–338.

16. Sorokin DY, Tourova TP, Kovaleva OL, Kuenen JG, Muyzer G. Aerobic carboxydotrophy under extremely haloalkaline conditions in *Alkalispirillum*/*Alkalilimnicola* strains isolated from soda lakes. Microbiology 2010; 156: 819–827.

17. Cypionka H, Meyer O, Schlegel HG. Physiological characteristics of various species of strains of carboxydobacteria. Arch Microbiol 1980; 127: 301–307.

18. King GM. Uptake of carbon monoxide and hydrogen at environmentally relevant concentrations by Mycobacteria. Appl Environ Microbiol 2003; 69: 7266–7272.

19. Gadkari D, Schricker K, Acker G, Kroppenstedt RM, Meyer O. *Streptomyces thermoautotrophicus* sp. nov., a thermophilic CO-and H_2_-oxidizing obligate chemolithoautotroph. Appl Environ Microbiol 1990; 56: 3727–3734.

20. O’Donnell AG, Falconer C, Goodfellow M, Ward AC, Williams E. Biosystematics and diversity amongst novel carboxydotrophic actinomycetes. Antonie Van Leeuwenhoek 1993; 64: 325–340.

21. Kim SB, Falconer C, Williams E, Goodfellow M. *Streptomyces thermocarboxydovorans* sp. nov. and *Streptomyces thermocarboxydus* sp. nov., two moderately thermophilic carboxydotrophic species from soil. Int J Syst Evol Microbiol 1998; 48: 59–68.

22. Krüger B, Meyer O. Thermophilic bacilli growing with carbon monoxide. Arch Microbiol 1984; 139: 402–408.

23. Kraut M, Meyer O. Plasmids in carboxydotrophic bacteria: physical and restriction analysis. Arch Microbiol 1988; 149: 540–546.

24. Dobbek H, Gremer L, Meyer O, Huber R. Crystal structure and mechanism of CO dehydrogenase, a molybdo iron-sulfur flavoprotein containing S-selanylcysteine. Proc Natl Acad Sci 1999; 96: 8884–8889.

25. Dobbek H, Gremer L, Kiefersauer R, Huber R, Meyer O. Catalysis at a dinuclear [CuSMo (O) OH] cluster in a CO dehydrogenase resolved at 1.1-Å resolution. Proc Natl Acad Sci 2002; 99: 15971–15976.

26. Meyer O, Schlegel HG. Biology of aerobic carbon monoxide-oxidizing bacteria. Annu Rev Microbiol 1983; 37: 277–310.

27. Conrad R, Meyer O, Seiler W. Role of carboxydobacteria in consumption of atmospheric carbon monoxide by soil. Appl Environ Microbiol 1981; 42: 211–215.

28. Conrad R. Soil microorganisms as controllers of atmospheric trace gases (H_2_, CO, CH_4_, OCS, N_2_O, and NO). Microbiol Mol Biol Rev 1996; 60: 609–640.

29. King GM. Molecular and culture-based analyses of aerobic carbon monoxide oxidizer diversity. Appl Environ Microbiol 2003; 69: 7257–7265.

30. Weber CF, King GM. Physiological, ecological, and phylogenetic characterization of Stappia, a marine CO-oxidizing bacterial genus. Appl Environ Microbiol 2007; 73: 1266–76.

31. Cunliffe M. Physiological and metabolic effects of carbon monoxide oxidation in the model marine bacterioplankton *Ruegeria pomeroyi* DSS-3. Appl Environ Microbiol 2013; 79: 738–740.

32. Cunliffe M. Correlating carbon monoxide oxidation with cox genes in the abundant marine *Roseobacter* clade. ISME J 2011; 5: 685.

33. King GM. Carbon monoxide as a metabolic energy source for extremely halophilic microbes: implications for microbial activity in Mars regolith. Proc Natl Acad Sci 2015; 4465–4470.

34. Hoeft SE, Blum JS, Stolz JF, Tabita FR, Witte B, King GM, et al. *Alkalilimnicola ehrlichii* sp. nov., a novel, arsenite-oxidizing haloalkaliphilic gammaproteobacterium capable of chemoautotrophic or heterotrophic growth with nitrate or oxygen as the electron acceptor. Int J Syst Evol Microbiol 2007; 57: 504–512.

35. Weber CF, King GM. The phylogenetic distribution and ecological role of carbon monoxide oxidation in the genus Burkholderia. FEMS Microbiol Ecol 2012; 79: 167–175.

36. Weber CF, King GM. Volcanic soils as sources of novel CO-Oxidizing *Paraburkholderia* and *Burkholderia*: *Paraburkholderia hiiakae* sp. nov., *Paraburkholderia metrosideri* sp. nov., *Paraburkholderia paradisi* sp. nov., <i>Paraburkholderia peleae<. Front Microbiol 2017; 8: 207.

37. Bartholomew GW, Alexander M. Microbial metabolism of carbon monoxide in culture and in soil. Appl Environ Microbiol 1979; 37: 932–937.

38. Patrauchan MA, Miyazawa D, LeBlanc JC, Aiga C, Florizone C, Dosanjh M, et al. Proteomic analysis of survival of *Rhodococcus jostii* RHA1 during carbon starvation. Appl Environ Microbiol 2012; 78: 6714–6725.

39. Yano T, Yoshida N, Takagi H. Carbon monoxide utilization of an extremely oligotrophic bacterium, *Rhodococcus erythropolis* N9T-4. J Biosci Bioeng 2012; 114: 53–55.

40. Park SW, Hwang EH, Park H, Kim JA, Heo J, Lee KH, et al. Growth of mycobacteria on carbon monoxide and methanol. J Bacteriol 2003; 185: 142–147.

41. King CE. Diversity and activity of aerobic thermophilic carbon monoxide-oxidizing bacteria on Kilauea Volcano, Hawaii. 2013.

42. Wu D, Raymond J, Wu M, Chatterji S, Ren Q, Graham JE, et al. Complete genome sequence of the aerobic CO-oxidizing thermophile Thermomicrobium roseum. PLoS One 2009; 4: e4207.

43. King CE, King GM. *Thermomicrobium carboxidum* sp. nov., and *Thermorudis peleae* gen. nov., sp. nov., carbon monoxide-oxidizing bacteria isolated from geothermally heated biofilms. Int J Syst Evol Microbiol 2014; 64: 2586–2592.

44. Islam ZF, Cordero PRF, Feng J, Chen Y-J, Bay S, Gleadow RM, et al. Two Chloroflexi classes independently evolved the ability to persist on atmospheric hydrogen and carbon monoxide. ISME J 2019; in press.

45. King CE, King GM. Description of *Thermogemmatispora carboxidivorans* sp. nov., a carbon-monoxide-oxidizing member of the class Ktedonobacteria isolated from a geothermally heated biofilm, and analysis of carbon monoxide oxidation by members of the class Ktedonobacter. Int J Syst Evol Microbiol 2014; 64: 1244–1251.

46. Nishimura H, Nomura Y, Iwata E, Sato N, Sako Y. Purification and characterization of carbon monoxide dehydrogenase from the aerobic hyperthermophilic archaeon *Aeropyrum pernix*. Fish Sci 2010; 76: 999–1006.

47. Sokolova TG, Yakimov MM, Chernyh NA, Lun’kova EY, Kostrikina NA, Taranov EA, et al. Aerobic carbon monoxide oxidation in the course of growth of a hyperthermophilic archaeon, *Sulfolobus* sp. ETSY. Microbiology 2017; 86: 539–548.

48. McDuff S, King GM, Neupane S, Myers MR. Isolation and characterization of extremely halophilic CO-oxidizing Euryarchaeota from hypersaline cinders, sediments and soils and description of a novel CO oxidizer, *Haloferax namakaokahaiae* Mke2. 3T, sp. nov. FEMS Microbiol Ecol 2016; 92.

49. Quiza L, Lalonde I, Guertin C, Constant P. Land-use influences the distribution and activity of high affinity CO-oxidizing bacteria Associated to type I-coxL genotype in soil. Front Microbiol 2014; 5: 271.

50. Greening C, Constant P, Hards K, Morales SE, Oakeshott JG, Russell RJ, et al. Atmospheric hydrogen scavenging: from enzymes to ecosystems. Appl Environ Microbiol 2015; 81: 1190–1199.

51. Ji M, Greening C, Vanwonterghem I, Carere CR, Bay SK, Steen JA, et al. Atmospheric trace gases support primary production in Antarctic desert surface soil. Nature 2017; 552: 400–403.

52. Constant P, Chowdhury SP, Pratscher J, Conrad R. Streptomycetes contributing to atmospheric molecular hydrogen soil uptake are widespread and encode a putative high-affinity [NiFe]-hydrogenase. Environ Microbiol 2010; 12: 821–829.

53. Berney M, Cook GM. Unique flexibility in energy metabolism allows mycobacteria to combat starvation and hypoxia. PLoS One 2010; 5: e8614.

54. Greening C, Villas-Bôas SG, Robson JR, Berney M, Cook GM. The growth and survival of *Mycobacterium smegmatis* is enhanced by co-metabolism of atmospheric H_2_. PLoS One 2014; 9: e103034.

55. Berney M, Greening C, Conrad R, Jacobs WR, Cook GM. An obligately aerobic soil bacterium activates fermentative hydrogen production to survive reductive stress during hypoxia. Proc Natl Acad Sci U S A 2014; 111: 11479–11484.

56. Greening C, Carere CR, Rushton-Green R, Harold LK, Hards K, Taylor MC, et al. Persistence of the dominant soil phylum Acidobacteria by trace gas scavenging. Proc Natl Acad Sci U S A 2015; 112: 10497–10502.

57. Liot Q, Constant P. Breathing air to save energy – new insights into the ecophysiological role of high-affinity [NiFe]-hydrogenase in Streptomyces avermitilis. Microbiologyopen 2016; 5: 47–59.

58. Christie-Oleza JA, Fernandez B, Nogales B, Bosch R, Armengaud J. Proteomic insights into the lifestyle of an environmentally relevant marine bacterium. ISME J 2012; 6: 124.

59. Muthusamy S, Lundin D, Mamede Branca RM, Baltar F, González JM, Lehtiö J, et al. Comparative proteomics reveals signature metabolisms of exponentially growing and stationary phase marine bacteria. Environ Microbiol 2017; 19: 2301–2319.

60. King GM. Contributions of atmospheric CO and hydrogen uptake to microbial dynamics on recent Hawaiian volcanic deposits. Appl Environ Microbiol 2003; 69: 4067–4075.

61. King GM, Weber CF, Nanba K, Sato Y, Ohta H. Atmospheric CO and hydrogen uptake and CO oxidizer phylogeny for Miyake-jima, Japan Volcanic Deposits. Microbes Environ 2008; 23: 299–305.

62. King GM, Weber CF. Interactions between bacterial carbon monoxide and hydrogen consumption and plant development on recent volcanic deposits. ISME J 2008; 2: 195–203.

63. Greening C, Berney M, Hards K, Cook GM, Conrad R. A soil actinobacterium scavenges atmospheric H_2_ using two membrane-associated, oxygen-dependent [NiFe] hydrogenases. Proc Natl Acad Sci U S A 2014; 111: 4257–4261.

64. Delgado-Baquerizo M, Oliverio AM, Brewer TE, Benavent-González A, Eldridge DJ, Bardgett RD, et al. A global atlas of the dominant bacteria found in soil. Science 2018; 359: 320–325.

65. Walsh CM, Gebert MJ, Delgado-Baquerizo M, Maestre F, Fierer N. A global survey of mycobacterial diversity in soil. bioRxiv 2019; 562439.

66. Snapper SB, Melton RE, Mustafa S, Kieser T, Jacobs WRJ. Isolation and characterization of efficient plasmid transformation mutants of *Mycobacterium smegmatis*. Mol Microbiol 1990; 4: 1911–1919.

67. Hartmans S, De Bont JA,. Aerobic vinyl chloride metabolism in *Mycobacterium aurum* L1. Appl Environ Microbiol 1992; 58: 1220–1226.

68. Gebhard S, Tran SL, Cook GM. The Phn system of *Mycobacterium smegmatis*: a second high-affinity ABC-transporter for phosphate. Microbiology 2006; 152: 3453–3465.

69. Cox J, Mann M. MaxQuant enables high peptide identification rates, individualized ppb-range mass accuracies and proteome-wide protein quantification. Nat Biotechnol 2008; 26: 1367.

70. Tyanova S, Temu T, Sinitcyn P, Carlson A, Hein MY, Geiger T, et al. The Perseus computational platform for comprehensive analysis of (prote) omics data. Nat Methods 2016; 13: 731.

71. Smith PK etal, Krohn R Il, Hermanson GT, Mallia AK, Gartner FH, Provenzano Md, et al. Measurement of protein using bicinchoninic acid. Anal Biochem 1985; 150: 76–85.

72. Walker JM. Nondenaturing polyacrylamide gel electrophoresis of proteins. The protein protocols handbook. 2009. Springer, pp 171–176.

73. Pruitt KD, Tatusova T, Maglott DR. NCBI reference sequences (RefSeq): a curated non-redundant sequence database of genomes, transcripts and proteins. Nucleic Acids Res 2007; 35: D61–D65.

74. Altschul SF, Gish W, Miller W, Myers EW, Lipman DJ. Basic local alignment search tool. J Mol Biol 1990; 215: 403–410.

75. Kumar S, Stecher G, Tamura K. MEGA7: Molecular Evolutionary Genetics Analysis version 7.0 for bigger datasets. Mol Biol Evol 2016; msw054.

76. Markowitz VM, Chen I-MA, Palaniappan K, Chu K, Szeto E, Grechkin Y, et al. IMG: the integrated microbial genomes database and comparative analysis system. Nucleic Acids Res. 2012., 40: D115–22

77. Leinonen R, Sugawara H, Shumway M, Collaboration INSD. The sequence read archive. Nucleic Acids Res 2010; 39: D19–D21.

78. Patel RK, Jain M. NGS QC Toolkit: a toolkit for quality control of next generation sequencing data. PLoS One 2012; 7: e30619.

79. Kopylova E, Noé L, Touzet H. SortMeRNA: fast and accurate filtering of ribosomal RNAs in metatranscriptomic data. Bioinformatics 2012; 28: 3211–3217.

80. Buchfink B, Xie C, Huson DH. Fast and sensitive protein alignment using DIAMOND. Nat Methods 2014; 12: 59.

81. Søndergaard D, Pedersen CNS, Greening C. HydDB: a web tool for hydrogenase classification and analysis. Sci Rep 2016; 6: 34212.

82. Janssen PH. Identifying the dominant soil bacterial taxa in libraries of 16S rRNA and 16S rRNA genes. Appl Environ Microbiol 2006; 72: 1719–1728.

83. Vorobev A V, Baani M, Doronina N V, Brady AL, Liesack W, Dunfield PF, et al. *Methyloferula stellata* gen. nov., sp. nov., an acidophilic, obligately methanotrophic bacterium that possesses only a soluble methane monooxygenase. Int J Syst Evol Microbiol 2011; 61: 2456–2463.

84. Tveit AT, Hestnes AG, Robinson SL, Schintlmeister A, Dedysh SN, Jehmlich N, et al. Widespread soil bacterium that oxidizes atmospheric methane. Proc Natl Acad Sci 2019; 201817812.

85. Ehhalt DH, Rohrer F. The tropospheric cycle of H_2_: a critical review. Tellus B 2009; 61: 500–535.

86. Greening C, Biswas A, Carere CR, Jackson CJ, Taylor MC, Stott MB, et al. Genomic and metagenomic surveys of hydrogenase distribution indicate H2 is a widely utilised energy source for microbial growth and survival. ISME J 2016; 10: 761–777.

87. Weber CF, King GM. Water stress impacts on bacterial carbon monoxide oxidation on recent volcanic deposits. ISME J 2009; 3: 1325–1334.

88. King GM. Microbial carbon monoxide consumption in salt marsh sediments. FEMS Microbiol Ecol 2007; 59: 2–9.

89. Myers MR, King GM. Perchlorate-coupled carbon monoxide (CO) oxidation: evidence for a plausible microbe-mediated reaction in Martian brines. Front Microbiol 2017; 8: 2571.

90. Parks DH, Chuvochina M, Waite DW, Rinke C, Skarshewski A, Chaumeil P-A, et al. A standardized bacterial taxonomy based on genome phylogeny substantially revises the tree of life. Nat Biotechnol 2018; 36: 996–1004.

