## Supplementary material for "Carbon monoxide dehydrogenases enhance bacterial survival by oxidising atmospheric CO"

**Table S1. List of genera known to oxidize CO.** The primary literature referenced was used to determine whether each genus has been reported to oxidize CO, grow chemolithoautotrophically on CO (column 3), or oxidize CO at atmospheric concentrations (column 4). Also listed is the phylogenetic clade that the CoxL sequences are affiliated with (column 6) based on the phylogenetic trees shown in **Fig. 4a** and **Fig. S2**. Note that all taxonomic assignments are based on the genome taxonomy database [90] and hence may differ from those historically reported. In this regard, the carboxydotroph [*Streptomyces*] *thermautotrophicus* is now recognized as a distinct genus from *Streptomyces* and the former class Betaproteobacteria is now the order Betaproteobacteriales in the class Gammaproteobacteria.

| **Class (Phylum)** | **Genus** | **Growth on CO?** | **Ambient uptake?** | **Reference** | **Enzyme clade** |
| --- | --- | --- | --- | --- | --- |
| Actinobacteria (Actinobacteriota) | [*Streptomyces*] | + | + | [19, 21] | Mixed 2 |
|  | *Mycobacterium* | ? | + | [18, 40] | Actinobacterial |
|  | *Rhodococcus* | ? | ? | [37–39] | Actinobacterial |
| Alphaproteobacteria (Proteobacteria) | *Oligotropha* | + | - | [13, 27] | Proteobacterial |
|  | *Bradyrhizobium* | +/- | + | [12, 14, 27, 29] | Proteobacterial |
|  | *Carbophilus* | + | ? | [12] | Unsequenced |
|  | *Ruegeria* | - | + | [29, 31, 32] | Proteobacterial |
|  | *Labrenzia* | - | + | [29, 30] | Proteobacterial |
|  | *Mesorhizobium* | - | + | [29] | Proteobacterial |
|  | *Aminobacter* | - | + | [29] | Proteobacterial |
|  | *Roseobacter* | - | ? | [32] | Proteobacterial |
|  | *Roseovarius* | - | ? | [32] | Proteobacterial |
|  | *Dinoroseobacter* | - | ? | [32] | Proteobacterial |
| Bacilli (Firmicutes) | *Hydrogenibacillus* | + | - | [22] | Mixed 1 |
|  | *Alicyclobacillus* | ? | ? | [41] | Mixed 1 |
|  | *Brevibacillus* | ? | ? | [41] | Mixed 1 |
|  | *Geobacillus* | ? | ? | [41] | Mixed 1 |
|  | *Anoxybacillus* | ? | ? | [41] | Unsequenced |
| Chloroflexia (Chloroflexota) | *Thermomicrobium* | - | + | [42–44] | Mixed 1 |
|  | *Sphaerobacter* | ? | ? | [41] | Mixed 1 |
| Deinococci (Deinococcota) | *Meiothermus* | ? | ? | [41] | Mixed 1 |
|  | *Thermus* | ? | ? | [41] | Mixed 1 |
| Gammaproteobacteria (Proteobacteria) | *Hydrogenophaga* | + | - | [12, 13, 15, 27] | Proteobacterial |
|  | *Zavarzinia* | + | ? | [12, 17] | Proteobacterial |
|  | *Alkalispirillum* | + | ? | [16] | Proteobacterial |
|  | *Xanthomonas* | +/- | + | [29] | Unsequenced |
|  | *Alkalilimnicola* | +/- | + | [16, 33, 34] | Proteobacterial |
|  | *Burkholderia* | - | + | [29, 35, 36] | Proteobacterial |
|  | *Paraburkholderia* | - | + | [36] | Proteobacterial |
|  | *Stenotrophomonas* | - | ? | [29] | Unsequenced |
| Ktedonobacteria (Chloroflexota) | *Thermogemmatispora* | - | + | [44, 45] | Mixed 1 |
| Halobacteria (Halobacterota) | *Halorubrum* | ? | + | [33, 48] | Halobacterial |
|  | *Haloferax* | ? | + | [48] | Halobacterial |
|  | *Haloterrigena* | ? | + | [48] | Halobacterial |
|  | *Haloarcula* | ? | ? | [48] | Halobacterial |
|  | *Natronorubrum* | ? | + | [33, 48] | Halobacterial |
| Thermoprotei (Crenarchaeota) | *Sulfolobus* | ? | ? | [47] | Mixed 1 |
|  | *Aeropyrum* | ? | ? | [46] | Mixed 1 |

**Table S2 (xlsx). Summary of proteome data.** Results are shown for a shotgun proteomic experiment that compared relative protein content of three carbon-replete (mid-exponential phase) and three carbon-limited (mid-stationary phase) cultures of *Mycobacterium smegmatis*.

**Table S3 (xlsx). List of carbon monoxide dehydrogenase sequences retrieved in this work.** A total of 709 sequences were retrieved of the form I CO dehydrogenase catalytic subunit (CoxL). Results are provided in table and FASTA format.

**Table S4.** **Comparison of four methods to determine apparent kinetic parameters for CO oxidation for whole cells of *Mycobacterium smegmatis*.**

| **Method** | ***V*_max app_ CO (nmol min g_dw_^-1^)** | ***K*_m app_ CO (nM)** |
| --- | --- | --- |
| Nonlinear regression | 3.13 | 350 |
| Lineweaver-Burk plot | 2.97 | 310 |
| Hanes-Woolf plot | 3.08 | 339 |
| Eadie-Hofstee plot | 2.90 | 291 |
| Average | 3.02 | 323 |

**Table S5 (xlsx). Details of the metagenome and metatranscriptome samples analyzed in this work.**

**Table S6 (xlsx). Relative abundance of carbon monoxide dehydrogenase and hydrogenase sequences in the analyzed metagenome and metatranscriptome datasets.** The relative abundance of CO dehydrogenase large subunit gene is shown by clade. The relative abundance of the hydrogenase large subunit genes, divided by subgroup, is also shown for comparison.

**Table S7. List of bacterial strains and plasmids used in this work.**

| **Name** | **Description** | **Reference** |
| --- | --- | --- |
| ***Mycobacterium smegmatis*** | |  |
| mc^2^155 | *ept-1*, efficient plasmid transformation mutant of mc^2^6 | [66] |
| Δ*coxL* | mc^2^155 with markerless deletion in *coxL* gene (MSMEG_0746) | This work |
| ***Escherichia coli*** | |  |
| TOP10 | F- *mcrA* Δ( *mrr-hsd*RMS-*mcr*BC) Φ80*lac*ZΔM15 Δ *lac*X74 *rec*A1 *ara*D139 Δ( *araleu*)7697 *gal*U *gal*K *rps*L (StrR) *end*A1 *nup*G | Thermo Fisher |
| ***Plasmids*** |  |  |
| pX33 | Gm^r^, *sacB*, mycobacterial Ts *ori*, p15A *ori*, *xylE* | [68] |
| pX33_coxL | 2245 bp fragment of left and right flanks of *coxL* gene (MSMEG_0746) in pX33 | This work |

**Table S8. List of primers used in this work.**

| **Purpose** | **Primer name** | **Sequence** | **Tm (°C)** |
| --- | --- | --- | --- |
| Sequencing of pX33_coxL to verify cloning of insert | Forward | TAATACGACTCACTATAGGG | 52.0 |
|  | Reverse | AATAGATCATCGTCGCCG | 51.9 |
| Screening of Δ*coxL*  (MSMEG_0746) mutant | Forward | CTGCCTATTGACGCTCGCG | 58.9 |
|  | Reverse | ACCACATCGTCGACACCGC | 60.4 |
| qRT-PCR of *coxL*  gene (MSMEG_0746) | Forward | CGTGGTGGTCAAACAGGAGA | 57.2 |
|  | Reverse | GATCTCGCCGACCATGATGT | 57.1 |
| qRT-PCR of *sigA*  gene (MSMEG_2758) | Forward | CTCAACGCCGAAGAAGAGGT | 57.2 |
|  | Reverse | GCCCTTGGTGTAGTCGAACT | 57.0 |

**Figure S1. Deletion of the *coxL* gene in *Mycobacterium smegmatis*.** The schematic shows the four main steps that led to the production of a knockout of the *coxL* gene (MSMEG_0746). (**i)** Construction of the pX33_coxL vector containing a fused left flank (LF) and right flank (RF) of the *coxL* gene. (**ii**) Temperature-mediated integration of the vector into the *M. smegmatis* chromosome *via* either the left flank or right flank of the chromosomal *coxL* gene. (**iii**) Chromosomal excision of the vector due to *sacB*-mediated sucrose toxicity to either wild-type revertants or Δ*coxL* mutants. (**iv**) PCR-based screening through primers targeting the flanks to confirm whether colonies are wild-type revertants (4610 bp product) or Δ*coxL* mutants (2330 bp product).

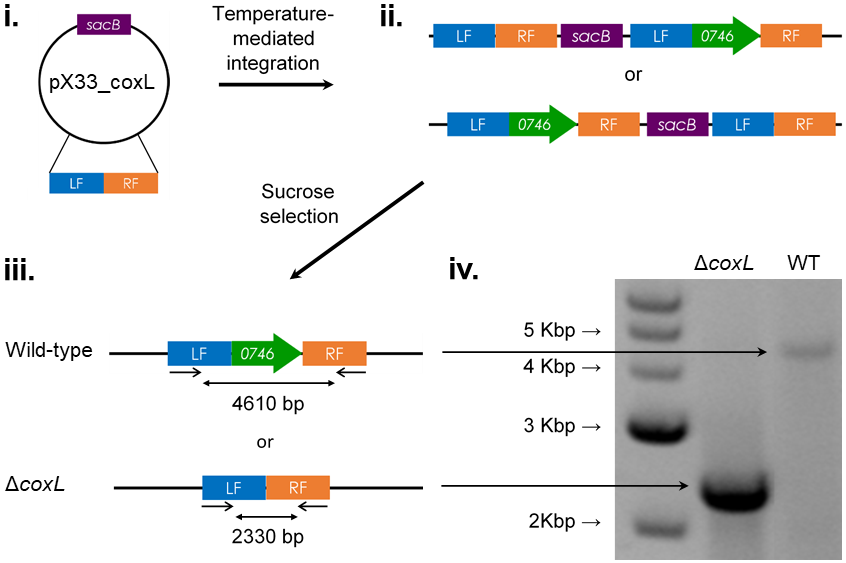

**Figure S2 (tif). Neighbor-joining** **phylogenetic tree showing the evolutionary history of the catalytic subunit of the form I carbon monoxide dehydrogenase (CoxL).** Evolutionary distances were computed using the Poisson correction model, gaps were treated by partial deletion, and the tree was bootstrapped with 500 replicates. The tree was constructed using all 709 CoxL sequences retrieved in this study as shown in **Table S3**. The tree is divided into five clades as per the maximum-likelihood tree **(Fig. 4a)**: Proteobacterial Clade (**a**), Mixed Clade 1 (**b**), Halobacterial Clade (**c**), Mixed Clade 2 (**d**), and Actinobacterial Clade (**e**). The tree was rooted with five form II CO dehydrogenase sequences (not shown). As the figure is too large to be embedded, it should be directly downloaded as a .tif file.

**Figure S3. Mirror diagram showing the abundance of genes and transcripts encoding the carbon monoxide dehydrogenase large subunit (*coxL*) by ecosystem type.** In total, 40 pairs of metagenomes and metatranscriptomes (20 aquatic, 20 terrestrial) were analyzed from a wide range of biomes (detailed in **Table S5**). The abundance of *hhyL* genes and transcripts, encoding the high-affinity group 1h [NiFe]-hydrogenase, are shown for comparison.

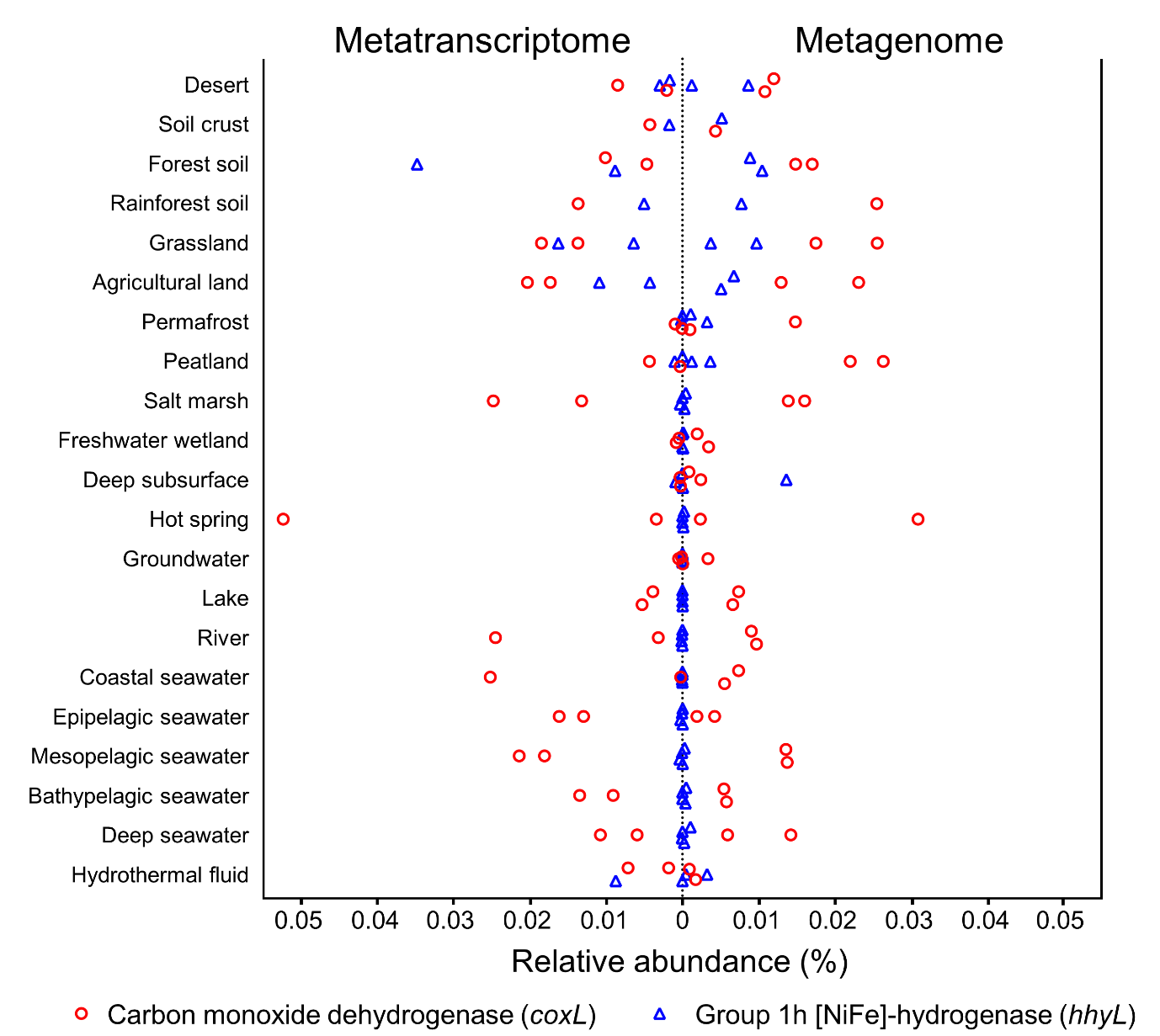
