## Supplementary figures and images for "Carbon monoxide dehydrogenases enhance bacterial survival by oxidising atmospheric CO"

### Figure S2

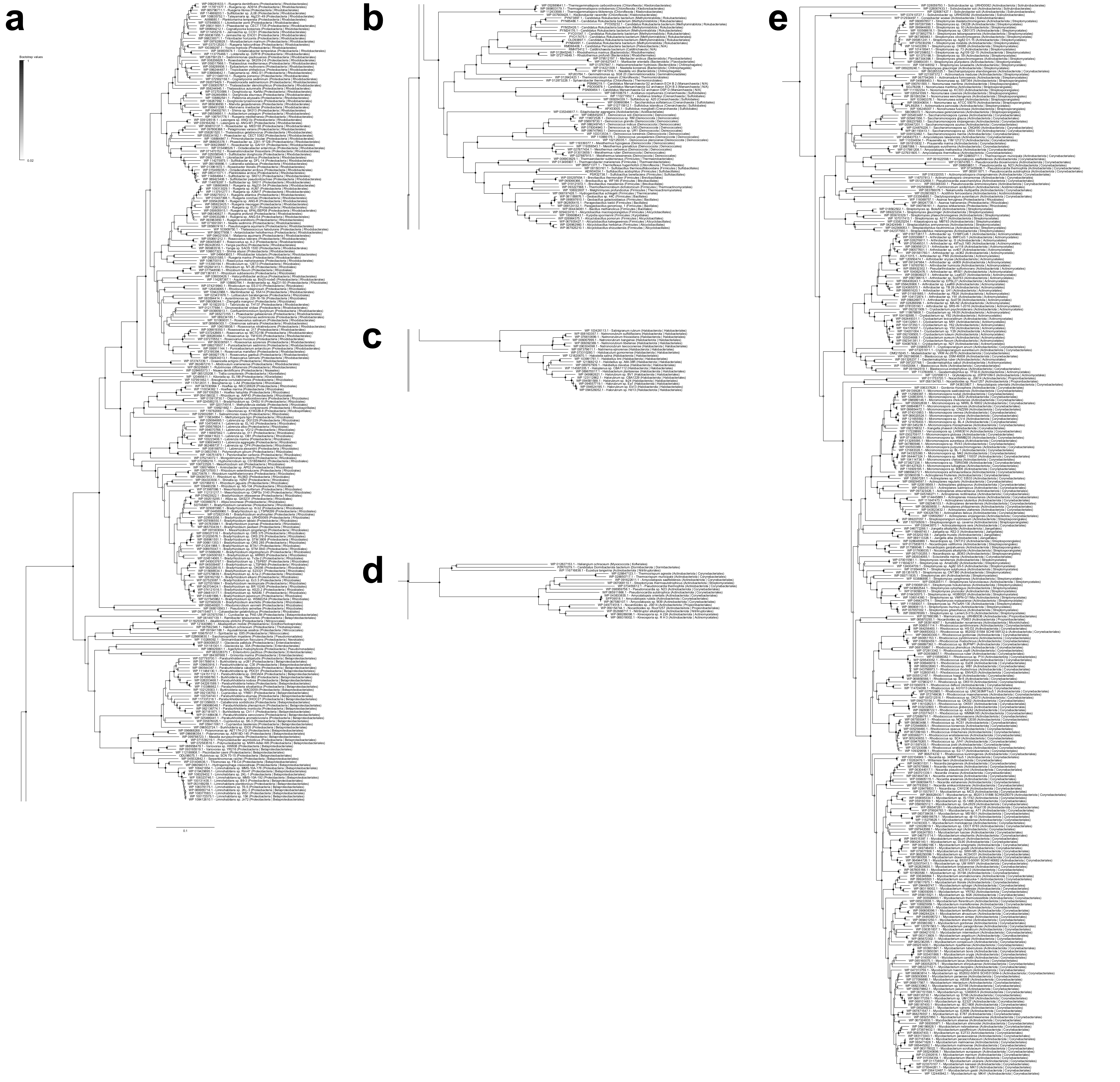
